# Carbon availability acts via cytokinins to promote gemma cup formation in *Marchantia polymorpha*

**DOI:** 10.64898/2025.12.08.692956

**Authors:** Jazmine L. Humphreys, Tom J. Fisher, Thibaut A. Perez, Eduardo Flores-Sandoval, Alessandro Silvestri, Ignacio Rubio-Somoza, Francois F. Barbier

## Abstract

Liverworts can clonally propagate by producing compact shoot structures called gemmae, which develop within basket-like structures known as gemma cups. It was previously reported in *Marchantia nepalensis* that carbon availability promotes gemma cup formation. However, the mechanisms by which carbon availability controls this process remains largely unexplored. To address this knowledge gap, we investigated how carbon promotes gemma cup formation using *Marchantia polymorpha* as a model species. Through a series of pharmacological and genetic experiments, we found that carbon availability promotes gemma cup formation by inducing the cytokinin pathway, thereby increasing the expression of *MpGCAM1* and *MpSTG*, which encode two transcription factors involved in forming the basal floor of gemma cups. Indeed, our data show that cytokinins accumulate in marchantia thallus in response to sucrose and to high light treatments. In addition, constitutive induction of cytokinin signalling could overcome the repressive effect of low sucrose on gemma cup formation, whereas suppression of this hormonal pathway led to inhibition of sucrose-induced gemma cup formation. Furthermore, our results indicate that sucrose can induce gemma cup formation independently of KAI2A and MAX2, two molecular components of karrikin signalling known to control this developmental process by inducing cytokinin synthesis. Interestingly, in flowering plants, carbon availability also promotes cytokinin accumulation to induce axillary bud outgrowth, a process involving the transcription factors AtRAX and AtLOF1, the *Arabidopsis thaliana* orthologues of MpGCAM1 and MpSTG, respectively. Collectively, these observations indicate that the interactions between carbon and cytokinins are critical for the developmental plasticity of land plants in response to their environment.

## Introduction

Land plants propagate by spreading spores or seeds that are produced through sexual reproduction. Certain species, such as *Selaginella moellendorffii*, *Hyperzia lucidula*, *Vittaria appaliachiana*, *Drosera scorpioides* and *Marchantia polymorpha* can clonally propagate by producing and dispersing propagules, also known as gemmae, which are compact shoots that stay dormant until they detach from the parent plant usually upon mechanical stimulation. Most liverworts, including *Marchantia polymorpha*, produce gemmae within specialised basket-like structures termed gemma cups, which develop on the dorsal surface of the thallus, the shoot portion of this type of plant (Kato et al., 2020).

Key genetic components regulating gemma cup formation have been identified, including the R2R3-MYB transcription factor GEMMAE CUP-ASSOCIATED MYB1 (MpGCAM1), which is required for the initiation of the basal floor during gemma cup development (Yasui et al., 2019). *MpGCAM1* expression is induced by cytokinin signalling (Aki et al., 2019a; Aki et al., 2022), which is triggered by an endogenous karrikin-like signal perceived by the KARRIKIN INSENSITIVE2A (MpKAI2A) receptor (Mizuno et al., 2021; Komatsu et al., 2023). MpKAI2A promotes cytokinin biosynthesis through LONELY GUY (MpLOG) (Komatsu et al., 2025), which activates cytokinin signalling through the two-component phosphorelay system and the MpRRB response regulator (Aki et al., 2019a; Aki et al., 2019b). This cytokinin-dependent pathway ultimately induces *MpGCAM1* expression, which confers stem cell-like properties to dorsal epidermal cells, and suppresses their default differentiation into air chamber tissue, thereby maintaining undifferentiated floor cells necessary for gemma initiation (Aki et al., 2022). Additional regulatory factors including SHOTGLASS (MpSTG), another R2R3-MYB transcription factor, play specialised architectural roles by maintaining *MpGCAM1* expression in the basal floor region while simultaneously suppressing air chamber development (Sakai et al., 2025). Post-transcriptional regulation via miR319 targeting of *MpRKD* (an RWP-RK domain transcription factor) further controls the spatial positioning and frequency of gemma cup formation along the thallus midrib (Sakai et al., 2025). Together, these factors coordinate the suppression of default tissue differentiation programmes while maintaining the stem cell-like state necessary for gemma cup morphogenesis (Sakai et al., 2025).

In addition to this genetic control, gemma cup formation is regulated by environmental cues. For example, in *Marchantia polymorpha*, gemma cup density is increased under short photoperiod compared to long photoperiod (Voth and Hamner, 1940). This developmental process is also controlled by mineral nutrition, and it was shown in the same species that gemma cup density increases with nitrate supply but decreases with phosphate supply (Voth and Hamner, 1940; Voth, 1941). It was reported in *Marchantia nepalensis* that higher light intensity and higher sucrose concentration promote gemma cup formation (Chopra and Sood, 1970), indicating plasticity of this developmental process in response to carbon. However, how carbon availability controls this process remains largely unexplored.

Carbon is the most abundant element in plants, representing about 35-40% of their dry matter. Plants obtain this element from atmospheric carbon dioxide (CO_2_) through photosynthesis, fixing it into organic molecules such as sugars. These sugars provide the carbon skeleton and energy required for growth and development. In contrast to other elements such as nitrogen or phosphate, plants cannot cope well with large variations in their carbon content, even in response to fluctuations in atmospheric CO_2_ levels (Cassan et al., 2024). Hence, plants must adjust their development to adapt to fluctuations in carbon availability and maintain constant levels of this element. The control of gemma cup formation by carbon availability in marchantia is a perfect example of this plasticity.

Shoot branching in angiosperms (flowering plants) is another good example of developmental plasticity in response to carbon availability. Indeed, the number of branches produced by a plant greatly depends on carbon availability modulated through light intensity or atmospheric CO_2_ concentration (Barbier et al., 2015b; Zhou et al., 2021; Fichtner et al., 2022; Patil et al., 2022; Swiegers et al., 2022). Shoot branching in angiosperms is driven by the outgrowth of axillary buds at the axils of leaves. The initiation of a new branch also depends on the balance of source-sink relationships among growing organs, reflecting the plant’s capacity to allocate carbon where it is most needed (Mason et al., 2014; Barbier et al., 2019b). Interestingly, recent studies have identified molecular components that are common to the formation of axillary buds in angiosperms and gemma cup formation in liverworts (Yasui et al., 2019; Kato et al., 2020; Sakai et al., 2025). Although not strictly comparable in terms of biological meaning, these developmental processes lead to the formation of “clonal shoots” on the main shoot of a plant.

Sugars do not simply regulate development through their trophic role. Indeed, several molecular mechanisms enable sugars to be sensed and to generate and transduce signals that regulate plant metabolism, physiology and ultimately development (Fichtner et al., 2021b; Li et al., 2021). Some of these sugar signalling pathways can interact with hormone signalling pathways to fine-tune plant development (Fichtner et al., 2021b; Mishra et al., 2022; Barbier et al., 2023; Wang et al., 2025). It was notably shown that the impact of sugar availability on shoot branching is mediated by signalling components such as HEXOKINASE1 or trehalose 6-Phosphate (Fichtner et al., 2017; Barbier et al., 2021; Fichtner et al., 2021a). Furthermore, sugar availability was reported to promote cytokinin accumulation and inhibit strigolactone signalling to promote shoot branching in different species of flowering plants (Bertheloot et al., 2020; Salam et al., 2021; Patil et al., 2022; Jiang et al., 2025).

*In silico* analyses of genes and proteins present in different groups of the green lineage suggest that some of the molecular mechanisms that mediate sugar signalling are ancient (Fichtner et al., 2021b; Lepper et al., 2025). However, while the impact of carbon availability and sugar signalling on plant development is well documented in angiosperms, little is known in other groups of land plants. This study aims to fill this knowledge gap by investigating how carbon availability controls gemma cup formation in *M. polymorpha*. Beyond unravelling the molecular mechanisms that underpin the carbon control of gemma cup formation in liverworts, this study advances our understanding of the mechanisms that govern plant evolution. Indeed, liverworts and flowering plants belong to groups of land plants that diverged around 450 million years ago. Studying and understanding the molecular mechanisms that underlie their developmental plasticity in response to the environment allows us to draw conclusions about the origins and genetics of the mechanisms that drive plant adaptation and evolution.

## Results

### Carbon availability promotes gemma cup formation in Marchantia polymorpha

Carbon availability was reported to control gemma cup formation in *Marchantia nepalensis* (Chopra and Sood, 1970). We first verified that this regulation also occurs in *Marchantia polymorpha*. To achieve this, we grew *M. polymorpha* for six weeks on plates supplemented with 5 or 50 mM sucrose. Under these conditions, gemma cup density was significantly higher under high sucrose than under low sucrose for both males (Mel-1) and females (Mel-2) of the Melbourne accession (Fig. 1A and B), confirming that carbon availability promotes gemma cup formation in this species.

**Figure 1.**
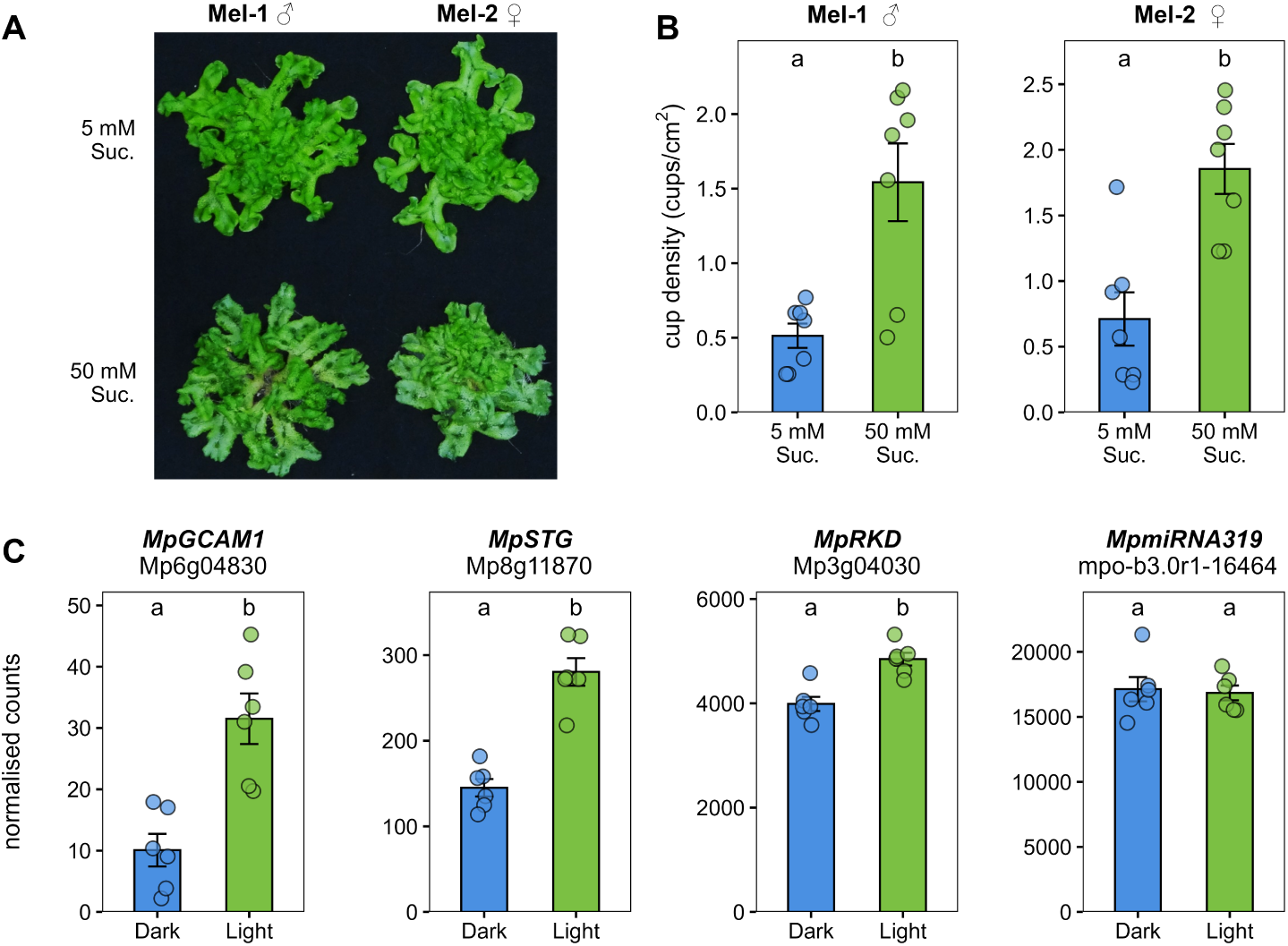
Carbon availability promotes gemma cup formation and induces expression of key regulatory genes in *Marchantia polymorpha*. (**A**) Representative images and (**B**) gemma cup density quantification of wild-type male (Mel-1, ♂) and female (Mel-2, ♀) thalli grown for six weeks on medium supplemented with 5 mM or 50 mM sucrose. (**C**) Expression of gemma cup regulatory genes in three-week-old thalli subjected to normal light (Light) or 6-h night extension (Dark). Normalised read counts are shown for *MpGCAM1* (Mp6g04830), *MpSTG* (Mp8g11870), *MpRKD* (Mp3g04030), and *MpmiRNA319* (mpo-b3.0r1-16464). Data represent mean ± SE (n = 6 biological replicates, shown as individual points). Different letters indicate statistically significant differences (P < 0.05, t-test for (**A**) and DESeq2 for (**C**)).

We next investigated whether carbon availability regulates key genes controlling gemma cup formation, including *MpGCAM1*, *MpSTG*, *MpRKD* and *miR319,* which targets *MpRKD*. Night extension is commonly used to deplete endogenous sugars in plants (Smith and Stitt, 2007). Therefore, we subjected 3-week-old thalli to either a 6 hr night extension or normal light conditions and performed mRNA-seq and small RNA-seq analysis. Compared to the samples exposed to a night extension, thalli under normal light conditions show 3-fold and 2-fold increases in *MpGCAM1* and *MpSTG,* respectively. *MpRKD* expression was modestly but significantly upregulated under normal light, whereas *miR319* levels remain unchanged. These results indicate that carbon availability primarily promotes the expression of *MpGCAM1* and *MpSTG* during gemma cup development.

### Carbon promotes cytokinin accumulation and requires cytokinin signalling to promote gemma cup formation

To gain further insight into the mechanism through which carbon promotes gemma cup formation, we looked at the expression of genes encoding molecular components acting upstream of *MpGCAM1*. Since this transcription factor is controlled by cytokinin signalling (Aki et al., 2022), we looked at cytokinin genes using the same transcriptomic data set from Fig. 1C. Our results show that the light condition slightly but significantly induced the expression of the cytokinin synthesis genes *ISOPENTENYL TRANSFERASE* (*MpIPT*) and *LONELY GUY* (*MpLOG*), compared to the dark treatment (extended night) (Fig. 2A). The expression of degradation genes *CYTOKININ OXIDASE/DEHYDROGENASE 1* and *2* (*MpCKX1* and *MpCKX2*) was also upregulated by light but more strongly than the synthesis genes (Fig. 2A). In contrast, the expression of genes involved in cytokinin perception, including those encoding CHASE DOMAIN CONTAINING HISTIDINE KINASE 1 (MpCHK1), the cytokinin receptor and HISTIDINE-CONTAINING PHOSPHOTRANSFER PROTEIN (MpHPT), that phosphorylates downstream transcription factors, were down regulated by the light treatment (Fig. 2A). The expression of the negative regulator *RESPONSE REGULAR A* (*MpRRA*) was strongly inhibited by light while the expression of *RESPONSE REGULAR B* (*MpRRB*), a positive mediator of cytokinin signalling, is unchanged (Fig. 2A). From these data it is clear that sugar availability controls cytokinin synthesis, degradation, perception and signalling. These data indicate that carbon availability broadly regulates the cytokinin pathway. However, because the expression of both biosynthesis and degradation genes was upregulated, we could not predict from gene expression alone whether cytokinin levels increase under high carbon conditions.

**Figure 2.**
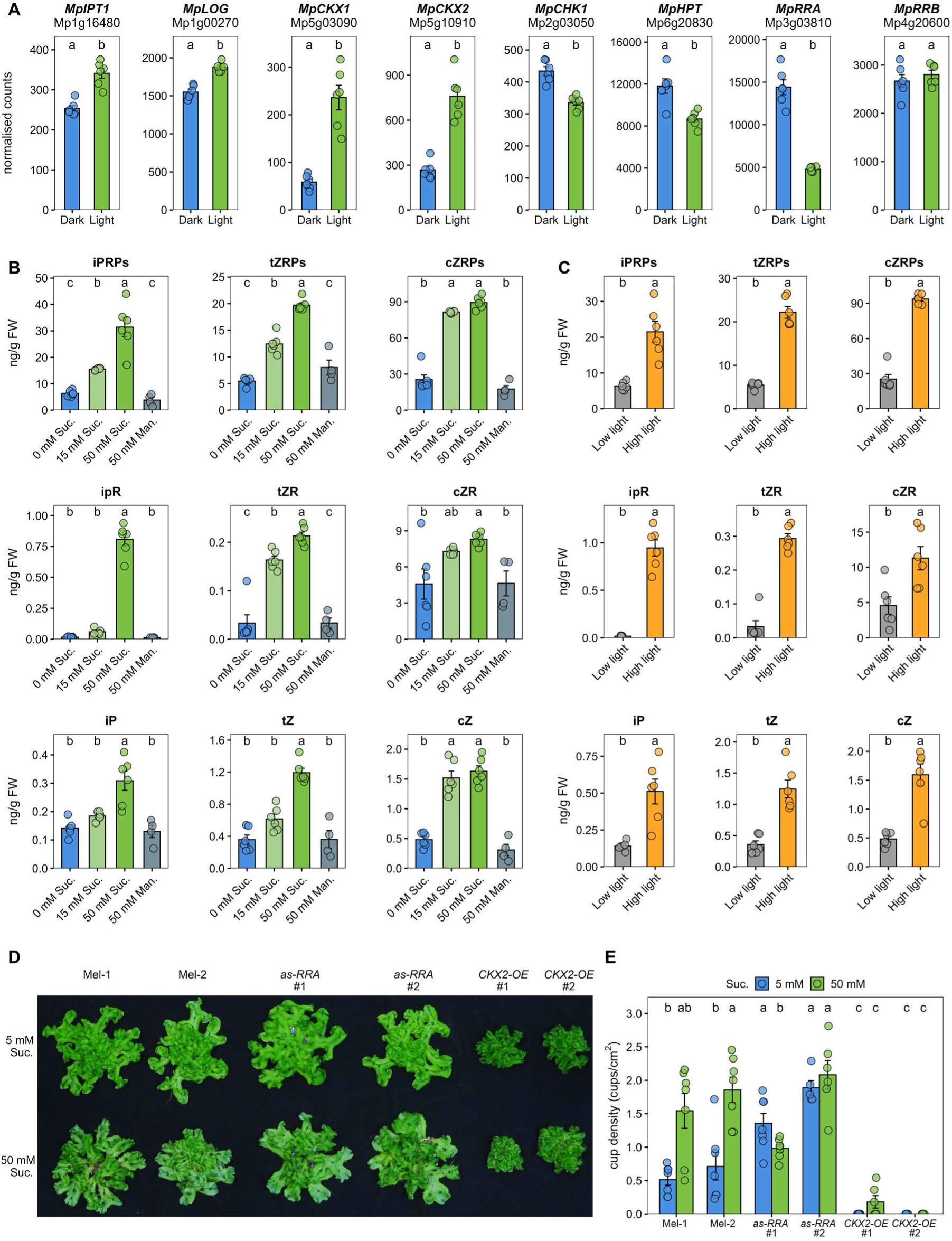
Carbon availability requires cytokinins to induce gemma cup formation. (**A**) Expression of cytokinin-related genes in three-week-old thalli subjected to normal light (Light) or 6-h night extension (Dark). (**B**) Cytokinin quantification by LC-MS/MS in thalli grown for three weeks on medium containing 0 mM, 15 mM, or 50 mM sucrose, or 50 mM mannitol as an osmotic control. (**C**) Cytokinin quantification in thalli grown for three weeks under low (40 µmol photons m⁻² s⁻¹) or high (110 µmol photons m⁻² s⁻¹) light. (**D**) Representative images and (**E**) gemma cup density quantification of wild-type (Mel-1, Mel-2), antisense *MpRRA* (*as-RRA* #1, #2), and *MpCKX2*-overexpressing (*CKX2-OE* #1, #2) thalli grown for six weeks on 5 mM or 50 mM sucrose. Data represent mean ± SE (n = 6 biological replicates, shown as individual points). Different letters indicate statistically significant differences (P < 0.05; DESeq2 in (**A**); one-way ANOVA with Tukey’s HSD post-hoc test in (**B**), (**C**), and (**E**)).

To directly assess whether carbon availability affects cytokinin accumulation in *M. polymorpha*, we quantified nine different isoforms of cytokinins in thallus tissues grown for three weeks on a range of sucrose concentrations (0, 15 and 50 mM). The results show that all nine isoforms, including the two active forms (iP and tZ), accumulated in a dose-dependent manner with sucrose concentration (Fig. 2B). Furthermore, cytokinins did not accumulate in response to 50 mM mannitol, a sugar used as an osmotic control (Fig. 2B), indicating that the cytokinin accumulation observed in response to 50 mM sucrose is not due to an osmotic effect.

To further support this conclusion, we modulated endogenous sugar levels using two light intensities (40 vs 110 µmol photons m⁻² s⁻¹). After three weeks of growth under these two light intensities, the quantification result clearly shows that all nine isoforms are upregulated by the higher light intensity. These observations further demonstrate that carbon availability increases cytokinin accumulation in *M. polymorpha*.

The results described above suggest that sugars act via cytokinins to promote gemma cup formation. To test this hypothesis, we generated transgenic lines that are altered in the cytokinin signalling pathway. We first created lines that mimic the presence of high levels of cytokinins by over-expressing an anti-sense construct that targets and silences *MpRRA* (*asRRA*), the negative cytokinin signalling component that is repressed by carbon availability (Fig. 2A), and grew them for 6 weeks on 5 and 50 mM sucrose. Overall, thallus morphology/ phenotype and cytokinin levels in asRRA lines were similar to wild-type plants (Fig. 2D). However, gemma cup quantification revealed that, unlike wild type plants (Mel-1 and Mel-2), which showed significantly fewer gemma cups at low sucrose concentration, two independent asRRA lines maintained high gemma cup density regardless of sucrose concentration (Fig. 2E). This result indicates that elevated cytokinin signalling is sufficient to overcome the inhibitory effect of low carbon availability on gemma cup formation.

We next disrupted the cytokinin pathway by creating two independent lines overexpressing CKX2 (*CKX2-OE*), which encodes a cytokinin degradation enzyme. These lines have very low levels of cytokinins compared to the WT plants (Supp. Fig. S1), and their thalli were small, compact, with densely ruffled, highly lobed margins, compared to the flat, expanded morphology of WT plants (Fig. 2D). Gemma cup density was extremely low in CKX2-OE lines, and increasing sucrose concentration did not alleviate this defect (Fig. 2E). Altogether, our observations suggest that the effect of sucrose on gemma cup induction could be mimicked by constitutively activating cytokinin signalling and was abolished when depleting cytokinins.

### Carbon availability promotes gemma cup formation independently of KAI2 signalling

The results above demonstrate that sugar availability promotes gemma cup formation through cytokinin signalling. It was previously reported that KAI2-dependent signalling, mediated by the F-box protein MORE AXILLARY GROWTH2 (MAX2), also controls gemma cup formation by inducing cytokinin biosynthesis through MpLOG (Komatsu et al., 2023; Komatsu et al., 2025). Moreover, studies in flowering plants showed that sugar availability can modulate MAX2-dependent strigolactone signalling to regulate shoot branching (Patil et al., 2022). We therefore investigated whether sugar availability acts through KAI2-MAX2-dependent signalling to regulate gemma cup formation in *M. polymorpha*.

We first looked at whether carbon availability regulates the expression of genes encoding MpKAI2A (the receptor), MpMAX2 (the F-box protein), and SUPRESSOR OF MAX2-LIKE (MpSMXL) (the transcriptional repressor targeted for degradation by the KAI2-MAX2 complex). In the night extension experiment, *MpMAX2* expression was modestly downregulated by light, while *MpKAI2A* and *MpSMXL* expression remained unchanged (Fig. 3A). These results suggest that carbon availability does not strongly regulate KAI2-ligand signalling at the transcriptional level.

**Figure 3.**
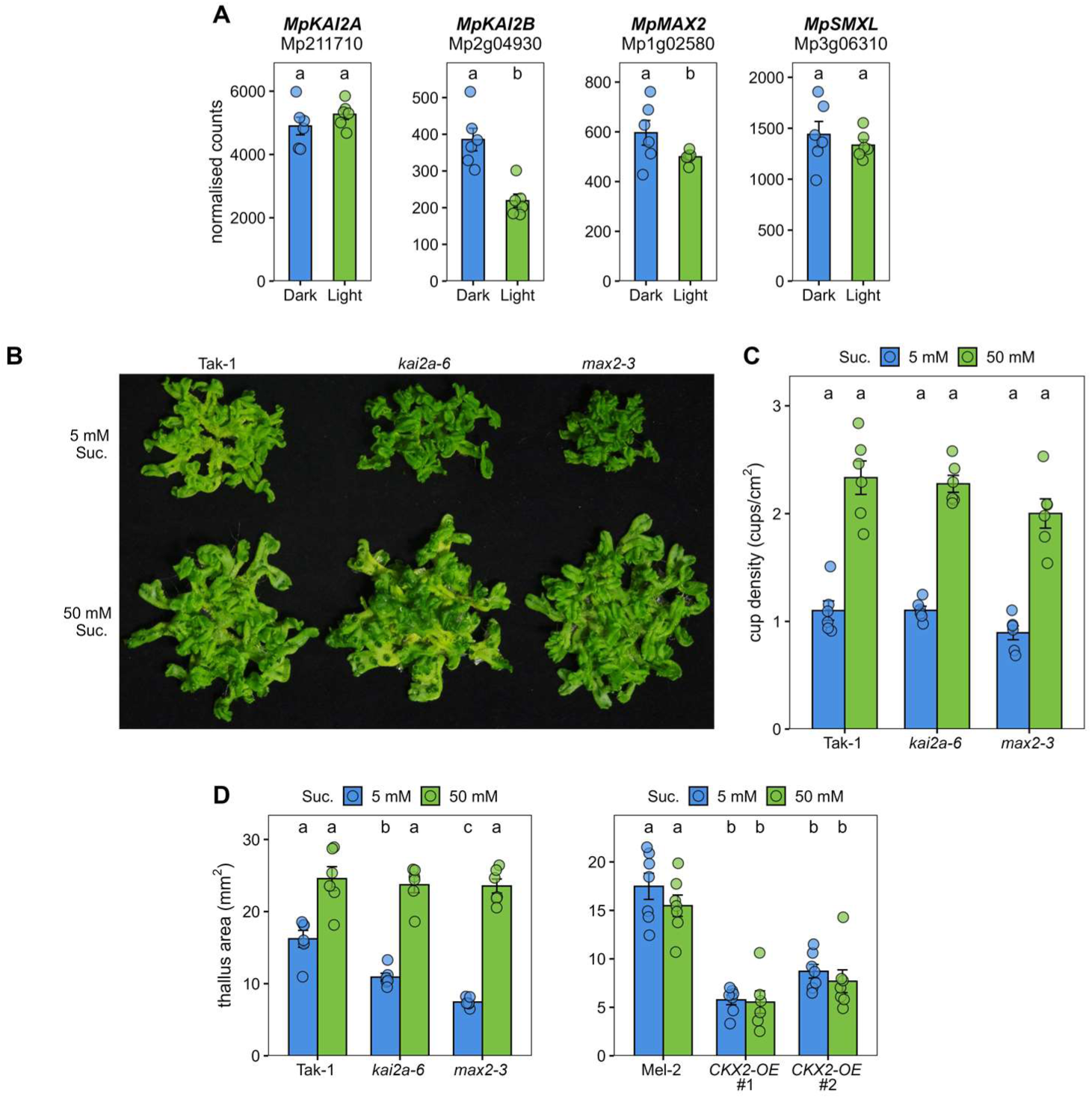
Carbon availability promotes gemma cup formation independently of KAI2 signalling. (**A**) Expression of KAI2-ligand signalling genes in three-week-old thalli subjected to normal light (Light) or 6-h night extension (Dark). (**B**) Representative images and (**C**) gemma cup density quantification of wild-type (Tak-1), *kai2a*-3, and *max2*-3 mutant thalli grown for six weeks on 5 mM or 50 mM sucrose. (**D**) Thallus area quantification of wild-type (Tak-1, Mel-2), *kai2a*-3, *max2*-3, and *CKX2-OE* (#1, #2) lines grown on 5 mM or 50 mM sucrose. Data represent mean ± SE (n = 6 biological replicates, shown as individual points). Different letters indicate statistically significant differences (P < 0.05; DEseq2 in (A); one-way ANOVA with Tukey’s HSD post-hoc test in (B) and (C)).

We next used a genetic approach to test whether carbon availability controls gemma cup formation through KL signalling. To achieve this we grew *kai2a-3* and *max2-3* mutants on 5 and 50 mM sucrose. Both mutants retained full sugar responsiveness, showing significantly increased gemma cup density at high sucrose comparable to wild-type controls (Fig. 3B and C). This demonstrates that sugar promotes gemma cup formation through a pathway independent of KAI2-MAX2 signalling.

Interestingly, *kai2a* and *max2* mutants grown on low sucrose resembled the cytokinin-depleted CKX-OE lines, showing reduced thallus size as quantified by projected area (Fig. 3D). This was expected, as these mutants produce reduced cytokinin levels (Komatsu et al., 2025). However, high sucrose concentration completely rescued this growth phenotype, restoring mutant thalli to wild-type appearance (Fig. 3B). Since sugar induces cytokinin accumulation in wild-type plants (Fig. 2B), we hypothesise that this sugar-responsive cytokinin biosynthesis compensates for the loss of KAI2-dependent cytokinin accumulation, explaining why these mutants retain normal development and gemma cup formation on high sugar medium.

## Discussion

In this study, we show that gemma cup formation in *Marchantia polymorpha* is promoted by carbon availability (Fig. 1A and B), consistent with previous observations in *Marchantia nepalensis* (Chopra and Sood, 1970). Our transcriptomic results indicate that carbon induces the expression of the two R2R3-MYB transcription factors *MpGCAM1* and *MpSTG* (Fig. 1C), which play critical roles in establishing and expanding the basal floor of the gemma cup (Yasui et al., 2019; Sakai et al., 2025). However, carbon availability did not strongly affect expression of *MpmiR319* or its target *MpRKD*, an RWP-RK domain transcription factor involved in the rim formation of the gemma cup (Sakai et al., 2025). These observations suggest that carbon availability promotes expression of components involved in basal floor establishment and expansion rather than those regulating rim formation (Fig. 4).

**Figure 4.**
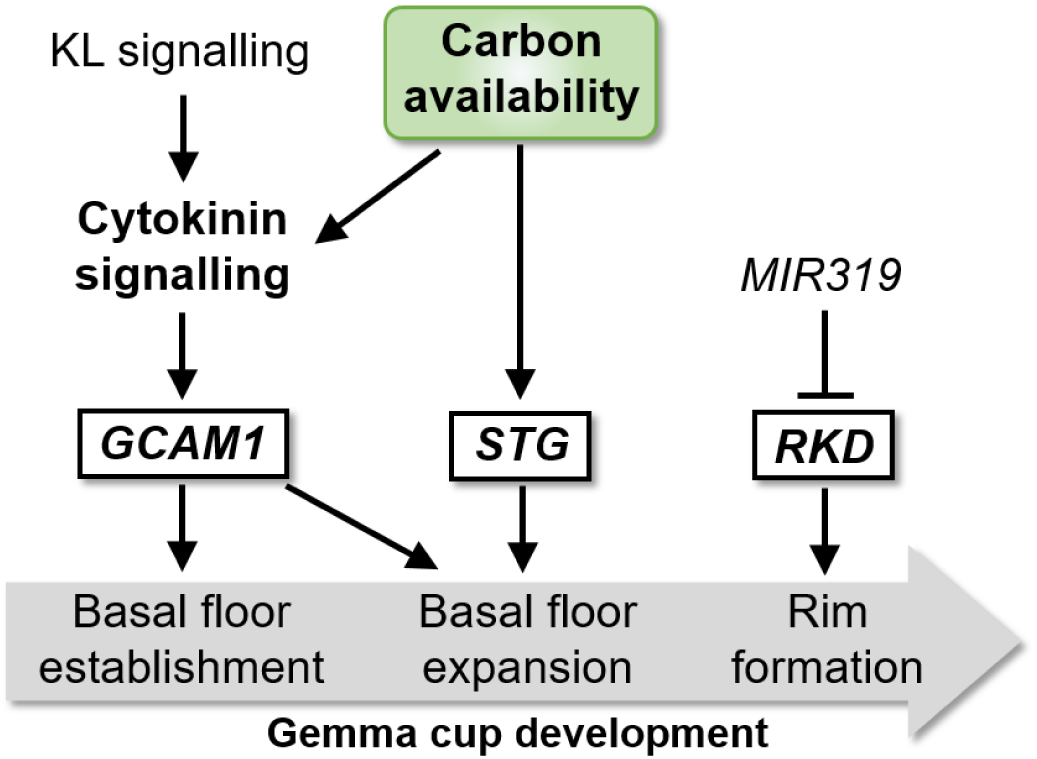
Proposed model of the involvement of carbon availability in the control of gemma cup formation in *Marchantia polymorpha* as described in the discussion. The outlined boxes represent the major transcription factors involved in gemma cup development. Adapted from Sakai *et al*., 2025.

Mass spectrometry quantification revealed that both high light intensity and sucrose treatment induced cytokinin accumulation in *M. polymorpha* thalli (Fig. 2B and C). Consistent with this, transcriptomic analysis showed that carbon availability upregulated cytokinin biosynthesis genes *MpIPT* and *MpLOG* (Fig. 2A), explaining the observed cytokinin accumulation. However, cytokinin degradation genes (*MpCKX1*, *MpCKX2*) were strongly upregulated, cytokinin perception genes (*MpCHK1*, *MpHPT*) were downregulated. We propose that this regulation is a negative feedback loop in response to the carbon-induced accumulation of cytokinins.

Genetic evidence demonstrated that cytokinins are required for sugar to promote gemma cup formation. Cytokinin-depleted CKX2-OE lines failed to form gemma cups even at high sucrose concentrations (Fig. 2D-E). Conversely, elevating cytokinin signalling by silencing the negative regulator MpRRA was sufficient to promote gemma cup formation under low sucrose concentration (Fig. 2D-E). Together, these results demonstrate that carbon availability promotes gemma cup formation by inducing cytokinin accumulation and signalling (Fig. 4).

The carbon-induced cytokinin biosynthesis observed in *M. polymorpha* is reminiscent of patterns reported in angiosperms, including *Arabidopsis thaliana*, *Rosa hybrida*, and *Solanum tuberosum* (Barbier et al., 2015a; Kiba et al., 2019; Salam et al., 2021; Jiang et al., 2025). Further transcriptomic analyses in *Arabidopsis thaliana* revealed that the expression of several *IPT* genes and other cytokinin-related genes is indeed upregulated by light using the same conditions as applied to marchantia in this study (Fig. S2). Interestingly, sugar-induced cytokinin biosynthesis has been implicated in promoting axillary bud outgrowth in angiosperms (Barbier et al., 2015a; Salam et al., 2021; Jiang et al., 2025). Interestingly, axillary bud formation in angiosperms is mediated by REGULATORS OF AXILLARY MERISTEMS (RAX) (Keller et al., 2006; Muller et al., 2006), which is orthologous to MpGCAM1 (Yasui et al., 2019). Our transcriptomic analysis revealed that *AtRAX3/MYB84* expression is also upregulated by light in *Arabidopsis thaliana* (Supp. Fig. S3), suggesting conserved carbon-responsive regulation of this developmental module. This observation is surprising because RAX controls axillary meristem initiation, a process generally considered to be genetically determined, in contrast to the well-documented environmental plasticity of axillary bud outgrowth (Barbier et al., 2019b).

These observations in bryophytes and angiosperms raise intriguing evolutionary questions. The convergence of sugar-cytokinin crosstalk regulating orthologous transcription factors (MpGCAM1/AtRAX) for lateral structure formation suggests two possible scenarios: either this regulatory module is ancestral, inherited from the common ancestor of bryophytes and angiosperms approximately 450 million years ago, or it arose independently through convergent evolution, reflecting a shared adaptive advantage of coupling carbon status to developmental plasticity. Distinguishing between these scenarios will require comparative analysis across early-diverging vascular plants and diverse bryophyte lineages to map the evolutionary trajectory of sugar-cytokinin interactions in plant development.

The KAI2-dependent signalling pathway promotes gemma cup formation by inducing the cytokinin pathway, notably by inducing the expression of the cytokinin synthesis gene *MpLOG* (Komatsu et al., 2025). Our results demonstrate that sucrose promotes this process independently of KAI2 and MAX2. Indeed, the expression of *MpMAX2* was downregulated by light while the expression of *MpKAI2A* was unchanged and those of *MpLOG* was upregulated (Fig. 2A and 3A), indicating that sucrose can induce cytokinin synthesis independently of the KAI2-signalling pathway. Furthermore, genetic analysis confirmed that *kai2a* and *max2* mutants retained full sugar responsiveness, showing increased gemma cup formation at high sucrose comparable to wild-type plants (Fig. 3B-C). This clearly indicates that carbon availability induces cytokinin -dependent gemma cup formation independently of KAI2-signalling (Fig. 4). Consistent with this model, high sucrose fully rescued the growth defects of *kai2a* and *max2* mutants (Fig. 3B,D). Because these mutants are cytokinin-deficient (Komatsu et al., 2025), and sugar induces cytokinin biosynthesis independently of KAI2-MAX2 signalling, this phenotypic rescue supports our conclusion that sugar-induced cytokinin biosynthesis can compensate for loss of KAI2-dependent cytokinin production. This is in line with a previous study showing that cytokinin treatment can rescue the decreased gemma cup formation of *kai2* and *max2* mutants (Komatsu et al., 2025).

In our experiments, *MpMAX2* expression was modestly downregulated by light (Fig. 3A), similar to patterns reported in angiosperms, albeit to a lesser extent (Barbier et al., 2015a; Patil et al., 2022), and as observed in our conditions in arabidopsis (Supp. Fig. S4). In this group of plants, the F-box protein MAX2 mediates both karrikin and strigolactone signalling, with strigolactones functioning as shoot branching inhibitors whose biosynthesis is strongly induced by phosphate and nitrate starvation (Gomez-Roldan et al., 2008; Umehara et al., 2010; Barbier et al., 2023; Waters and Nelson, 2023). Interestingly, phosphate starvation-induced upregulation of strigolactone biosynthesis, well-characterised in angiosperms, is also conserved in bryophytes including liverworts (Decker et al., 2017; Kodama et al., 2022), indicating that nutrient-hormone regulatory networks predate the bryophyte-angiosperm divergence. These observations suggest that the integration of nutrient sensing with hormonal regulation represents an ancient feature of land plant biology, not limited to carbon-cytokinin interactions. It is tempting to speculate that certain metabolites, such as cytokinins and strigolactones, were co-opted into signalling pathways because their nutrient-responsive regulation conferred adaptive advantages for responding to environmental variation. This integration may have been crucial during land colonisation, allowing early terrestrial plants to coordinate growth and development with resource availability in the challenging and heterogeneous terrestrial environment. Comparative analyses across bryophytes, early-diverging vascular plants, and charophyte algae are needed to reconstruct the evolutionary origins of these nutrient-hormone regulatory networks.

## Acknowledgment

We would like to thank Junko Kyozuka (Tohoku University, Japan) for providing genetic material, Steve Smith (University of Tasmania, Australia) for fruitful discussions, and Alejandro Correa-Lozano and David Nichols (CSL, University of Tasmania, Australia) for advice and assistance with hormone analysis. This work was supported by the Australian Research Council Centre of Excellence for Plant Success in Nature and Agriculture (ARC grant no. CE200100015).

## Material and methods

### Plant material and transgenic lines

For *Marchantia polymorpha*, all experiments were carried out in the Melbourne (Mel-1, male and Mel-2, female) Australian strain (Flores-Sandoval et al., 2015; Flores-Sandoval et al., 2025) or in the Takaragaike-1 (Tak-1, male) Japanese strain (Ishizaki et al., 2016) as indicated in the figure legends. The *kai2a-6* and *max2-3* mutants were provided by Junko Kyozuka and described before (Mizuno et al., 2021). The *asRRA* and *CXK2-OE* lines were created as described below. For *Arabidopsis thaliana*, all experiments were performed in the Columbia-0 (Col-0) accession.

### Cloning

The Mp*RRA* artificial microRNA (amiR) targets Mp*RRA*/Mp3g03810 mRNA based off a 21bp sequence (GAGCCACTGCGCCCCAGCTGA), which was modified to be incorporated into an amiR construct as described in (Flores-Sandoval et al., 2016). This amiR-MpRRA was synthesized (GeneScript) and subcloned into EcoRI-cut pENTR2B entry vector (Invitrogen, A10463) using a NEBuilder HiFi DNA Assembly Master Mix (New England Biolabs, E2621), with primers (pE2B)Mpmir160F (GTCGACTGGATCCGGTACCGGCACCTCCTCTCTCCGACTGC) and Mpmir160(pE2B)R (ATATCTCGAGTGCG GCCGCGTAAGTAAATCTATCAAACATC).

MpCKX2/Mp5g10910 coding sequence (CDS) was originally cloned from *Marchantia polymorpha* collected in Melbourne, Australia (Flores-Sandoval et al., 2015). This MpCKX2(CDS) was subcloned into EcoRI-cut pENTR2B entry vector (Invitrogen, A10463) using a NEBuilderHiFi DNA Assembly Kit (New England Biolabs, E2621), with primers ASSpE2b-MpCKX2-F1 (GTCGACTGGATCCGGTACCGATGATGCTGCAATTACTGAAATA) and ASSpE2b-MpCKX2-R1 (ATATCTCGAGTGCGGCCGCGTCATAATAGGAACGGGAATGTCT).

The destination vector pMpGWB337-tdtNLS (Nishihama et al., 2016; Flores-Sandoval et al., 2025) was recombined with amiR-MpRRA pENTR2B and MpCKX2 pENTR2B using LR clonase II (Invitrogen, 11791) to produce the binary vectors amiR-MpRRA GWB337 and MpCKX2 GWB337, respectively.

### Transformation and screening

Sporophytes were collected in the lab from a cross between Mel-1 and Mel-2. Transformation of *M. polymorpha* sporelings with the binary vectors amiR-MpRRA GWB337 for *asRRA* lines and MpCKX2 GWB337 for *CKX2-OE* lines followed the description by (Ishizaki et al., 2008), using sporeling liquid media of half strength B5 (½B5) (Phytotech Labs, G398) (Gamborg et al., 1968) supplemented with 2% sucrose (Merck), 0.1% casamino acids (Phytotech Labs, G229) and 0.03% l-glutamine (MP Biomedicals).

Sporelings were screened with a combination of 0.5μM chlorsulfuron in media to select for integration and expression of the GWB337 T-DNA, visual phenotyping aided by published mutant descriptions (Aki et al., 2019a) and checked for the absence of any tdtNLS fluorescent sectors, as this would indicate accidental Cre recombination and therefore loss of amiR-MpRRA or MpCKX2 CDS. Subsequently, each mutant line was purified of chimerism by propagation from a single gemma.

### Growth conditions and treatments

Marchantia plants were grown in growth cabinets with a photoperiod of 16 hours and a temperature of 19 ±1°C at night and 22 ±1°C during the day. Gemmae were placed in the centre of plates filled with half-strength Gamborg B5 + vitamins (Duchefa) with or without sucrose as indicated in the figure legends. In figure 2, mannitol was used as an osmotic control. Plants used for sugar feeding experiments were grown with a light intensity of 40 ±10 μmol m⁻² s⁻¹. Arabidopsis plants were grown in growth cabinets with a photoperiod of 16 hours and a temperature of 19 ±1°C at night and 22 ±1°C during the day and a light intensity of 120 ±10 μmol m⁻² s⁻¹.

For marchantia, the night extension was applied on three-week-old plants grown under 60 ±10 μmol m⁻² s⁻¹ with the same growth conditions as described above. At the beginning of the night, half of the plants were covered. Plants were harvested six hours after the light turned on for further analysis. For arabidopsis, plants were grown for three weeks under a 16 hr-photoperiod with a light intensity of 60 ±10 μmol m⁻² s⁻¹. The night extension was applied as described above.

For cytokinin analysis of sugar feeding experiments, three-week-old plants grown on 0, 15, or 50 mM sucrose, or 50 mM mannitol were harvested and immediately frozen in liquid nitrogen (six biological replicates per treatment). Additionally, plants grown for three weeks under low (40 µmol photons m⁻² s⁻¹) or high (110 µmol photons m⁻² s⁻¹) light intensity were harvested for cytokinin quantification (six biological replicates per condition).

### Gemma cup quantification

After six weeks of treatment, the number of gemma cups per plant was scored and the projected thallus surface was determined from scanned images using ImageJ. Cup density was determined by dividing the number of cups per plant by the thallus area.

### RNA extraction

Three-week-old *M. polymorpha* (Mel accession) and *A. thaliana* (Col-0) plants were subjected to either normal light conditions or a 6-hour night extension treatment as described above. Six biological replicates per condition were collected for each species. Total RNA was extracted from ground frozen tissue using the phenol–chloroform–free CTAB/PVP-based method previously described (Barbier et al., 2019a). Frozen samples were homogenised under liquid nitrogen and lysed in CTAB/PVP buffer supplemented with DTT. After clarification by centrifugation, RNA was precipitated from the supernatant with isopropanol and washed with ethanol. The pellet was resuspended in RNase-free water and treated with DNase I to remove residual genomic DNA, followed by a second round of isopropanol precipitation and ethanol washing. The purified RNA was quantified and assessed for integrity prior to downstream applications.

### RNA library preparation and sequencing

Total RNA was sent to the Ramaciotti Centre for Genomics, Sydney, Australia (www.ramaciotti.unsw.edu.au) for library preparation and sequencing. Libraries were sequenced on a NovaSeq 6000 generating 100bp paired end reads with a target depth of 25 million reads per sample. For small RNA-seq, libraries were prepared from the same RNA samples and sequenced on NovaSeq 6000 generating 75bp paired end reads with a target depth of 25 million reads per sample.

### mRNA seq analysis

Raw sequencing reads were assessed for quality using FastQC (v 0.12.1) (Andrews, 2010) Adapter sequences and low-quality bases were removed using Trimmomatic (v 0.39) (Bolger et al., 2014) with the following parameters: the first 10 nucleotides were trimmed from each read using HEADCROP, and a sliding window approach (SLIDINGWINDOW:4:20) was applied to remove bases with quality scores below 20. For *M. polymorpha*, quality-filtered reads were pseudo-aligned to the v7.1 reference transcriptome using kallisto (0.50.0) (Bray et al., 2016) with default parameters. For *A. thaliana*, reads were pseudo-aligned to the TAIR10 reference transcriptome. Transcript-level abundance estimates were imported into R and collapsed to gene-level counts using tximport (v1.36.0) (Soneson et al., 2016) with a transcript-to-gene mapping file. Differential expression analysis was performed using DESeq2 (v1.48.2) (Love et al., 2014) with default parameters, including independent filtering of low-count genes. Genes with an adjusted p-value (padj) < 0.05 were considered differentially expressed.

### Small RNA-seq analysis

Small RNA sequencing reads were quality-checked using FastQC (v0.11.5) and adapter/quality trimmed with Trim Galore (v0.4.5) using the parameters “-q 20 --length 16 --max_length 33”, which also generated post-trim FastQC reports. Trimmed reads were mapped to small RNA–generating loci annotated in the Plant Small RNA Genes database (https://plantsmallrnagenes.science.psu.edu/ genomes.php?genomes_id=49), which provides consistent annotation of sRNA loci across plant species (Lunardon et al., 2020). Read alignment was performed with HISAT2 (v2.1.0) using a single-end alignment strategy and allowing up to 100 reportable alignments per read (-k 100), followed by filtering to retain reads with zero mismatches and subsequent sorting using SAMtools (v1.15.1). Locus-level read quantification was carried out with HTSeq (v0.12.4) using htseq-count with settings “-s no -t exon --nonunique all -a 0 --secondary-alignments score”. Differential expression analysis of sRNA loci was performed using DESeq2 with default parameters. Small RNA loci with padj < 0.05 were considered differentially expressed. Annotation information including sRNA sequence, family classification (miRNA, siRNA, or other), and overlapping gene features was obtained from the Plant Small RNA Genes database.

### Hormone extraction and purification

Frozen tissue samples were weighed, transferred to 1.5 mL microcentrifuge tubes containing steel beads and immediately frozen in liquid nitrogen. Samples were homogenised using a TissueLyser (30 Hz for 1 minute) and extracted with 1 mL of 80% (v/v) methanol containing 250 mg/L butylated hydroxytoluene (BHT). To each sample, 100 µL of deuterated cytokinin internal standard mixture (10 ng per standard) containing ²H₅-trans-zeatin, ²H₅-trans-zeatin riboside, ²H₃-dihydrozeatin riboside, ²H₃-dihydrozeatin, ²H₆-isopentenyladenine, ²H₆-isopentenyladenosine, [¹⁵N₄]-cis-zeatin, [¹⁵N₄]-cis-zeatin riboside, and [²H₃]-dihydrozeatin riboside-5’-monophosphate sodium salt was added. Samples were incubated overnight at 4°C with gentle agitation. Following incubation, samples were centrifuged at 3,000 xg for 10 minutes at 4°C, and supernatants were transferred to glass scintillation vials. Samples were purified using solid-phase extraction (SPE) with C18 columns (Waters, Australia) preconditioned with 80% methanol. The eluate was collected and dried under nitrogen in a sample concentrator. Dried extracts were resuspended in 150 µL of 20% methanol, centrifuged at 14,000 xg for 3 minutes, and transferred to autosampler vials for LC-MS/MS analysis.

### LC-MS/MS analysis

Cytokinin analysis was performed using a Waters Acquity H-Class UPLC coupled to a Waters Xevo triple quadrupole mass spectrometer. Nine cytokinin forms were quantified: the free bases N⁶-isopentenyl adenine (iP), trans-zeatin (tZ), and cis-zeatin (cZ); the ribosides isopentenyladenosine (iPR), trans-zeatin riboside (tZR), and cis-zeatin riboside (cZR); and the ribotides (5’-monophosphates) isopentenyl adenosine-5’-monophosphate (iPRP), trans-zeatin riboside-5’-monophosphate (tZRP), and cis-zeatin riboside-5’-monophosphate (cZRP). Cytokinin concentrations were calculated using the isotope dilution method: Cytokinin level (ng/g FW) = (peak area endogenous × ng standard added) / (peak area standard × tissue fresh weight in g)

### Statistical analysis

For hormone measurements and gemma cup phenotyping, statistical analyses were performed in R (v4.5.1) Differences between treatments were assessed using one-way analysis of variance (ANOVA) followed by Tukey’s Honest Significant Difference (HSD) post-hoc test. Significance was set at p < 0.05. Data are presented as mean ± standard error.

## Supplementary Figures

**Supplementary Figure S1.**
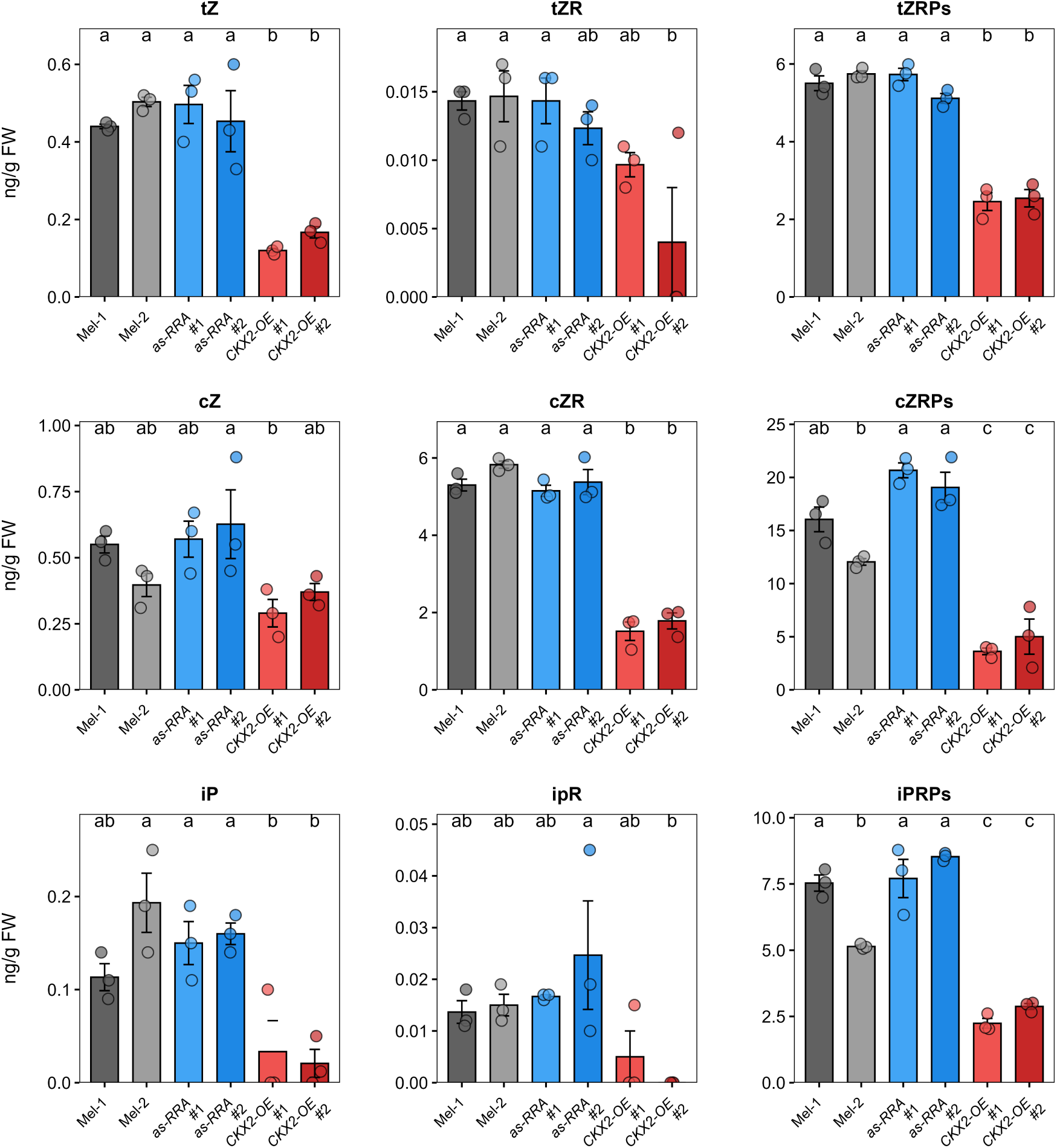
Cytokinin levels in wild-type and cytokinin signalling lines. Quantification of cytokinin isoforms by LC-MS/MS in wild-type (Mel-1, Mel-2), antisense MpRRA (*as-RRA* #1, #2), and MpCKX2-overexpressing (*CKX2-OE* #1, #2) thalli. Cytokinin ribotide precursors (iPRPs, tZRPs, cZRPs), ribosides (ipR, tZR, cZR), and free bases (iP, tZ, cZ) are shown. Data represent mean ± SE (n = 3 biological replicates, shown as individual points). Different letters indicate statistically significant differences (P < 0.05, one-way ANOVA with Tukey’s HSD post-hoc test).

**Supplementary Figure S2.**
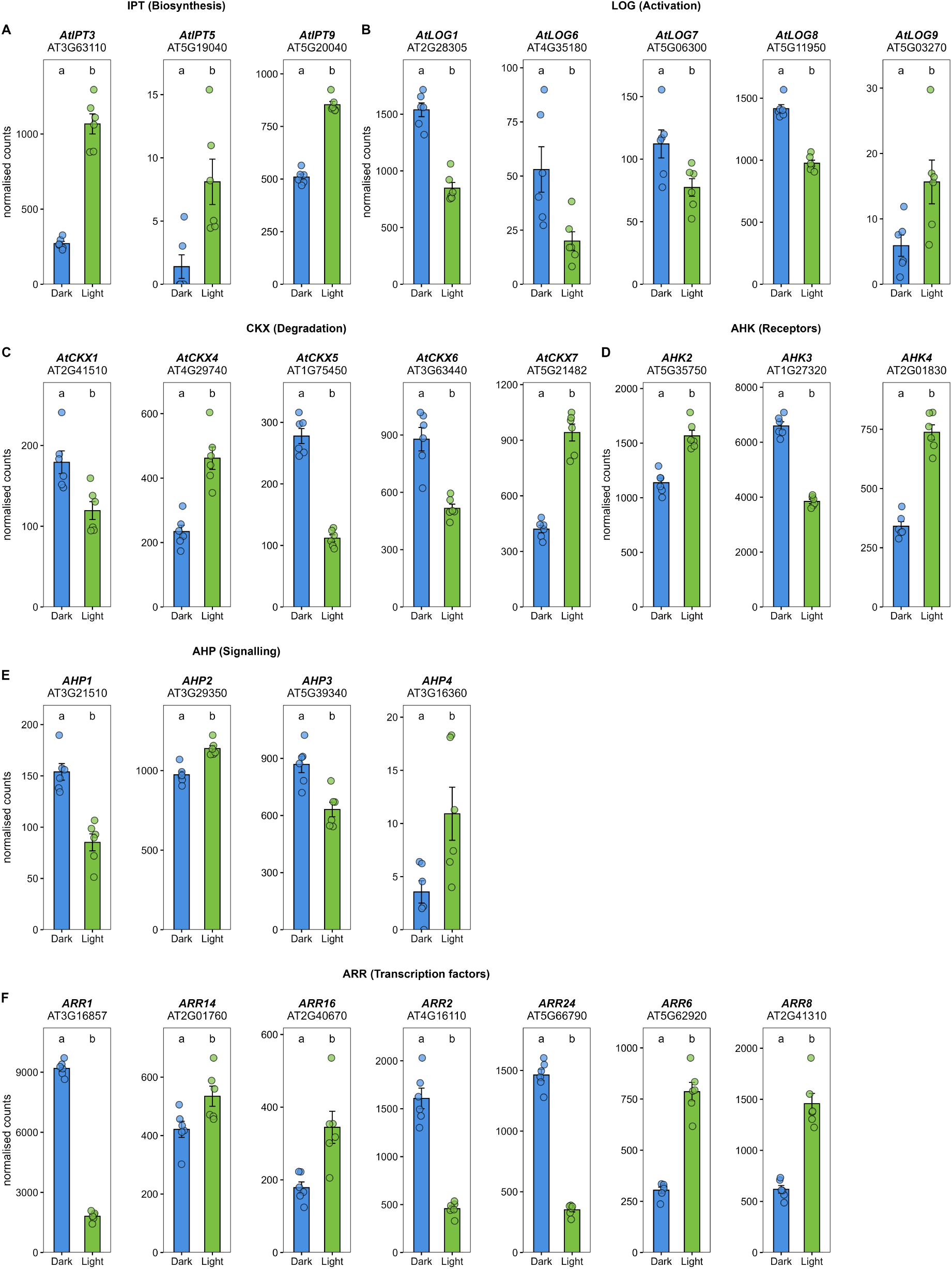
Impact of carbon availability on the expression of cytokinin genes in *Arabidopsis thaliana*. Expression of cytokinin genes in *Arabidopsis thaliana* three-week-old rosettes subjected to normal light (Light) or dark (Dark) conditions. Only genes showing significant differential expression are shown. (**A**) *IPT* genes (cytokinin biosynthesis). (**B**) *AHK* genes (cytokinin receptors). (**C**) *AHP* genes (histidine phosphotransfer). (**D**) *LOG* genes (cytokinin activation). (**E**) *CKX* genes (cytokinin degradation). (**F**) *ARR* genes (response regulators). Data represent mean ± SE of normalised read counts (n = 6 biological replicates, shown as individual points). Different letters indicate statistically significant differences (P < 0.05, DESeq2).

**Supplementary Figure S3.**
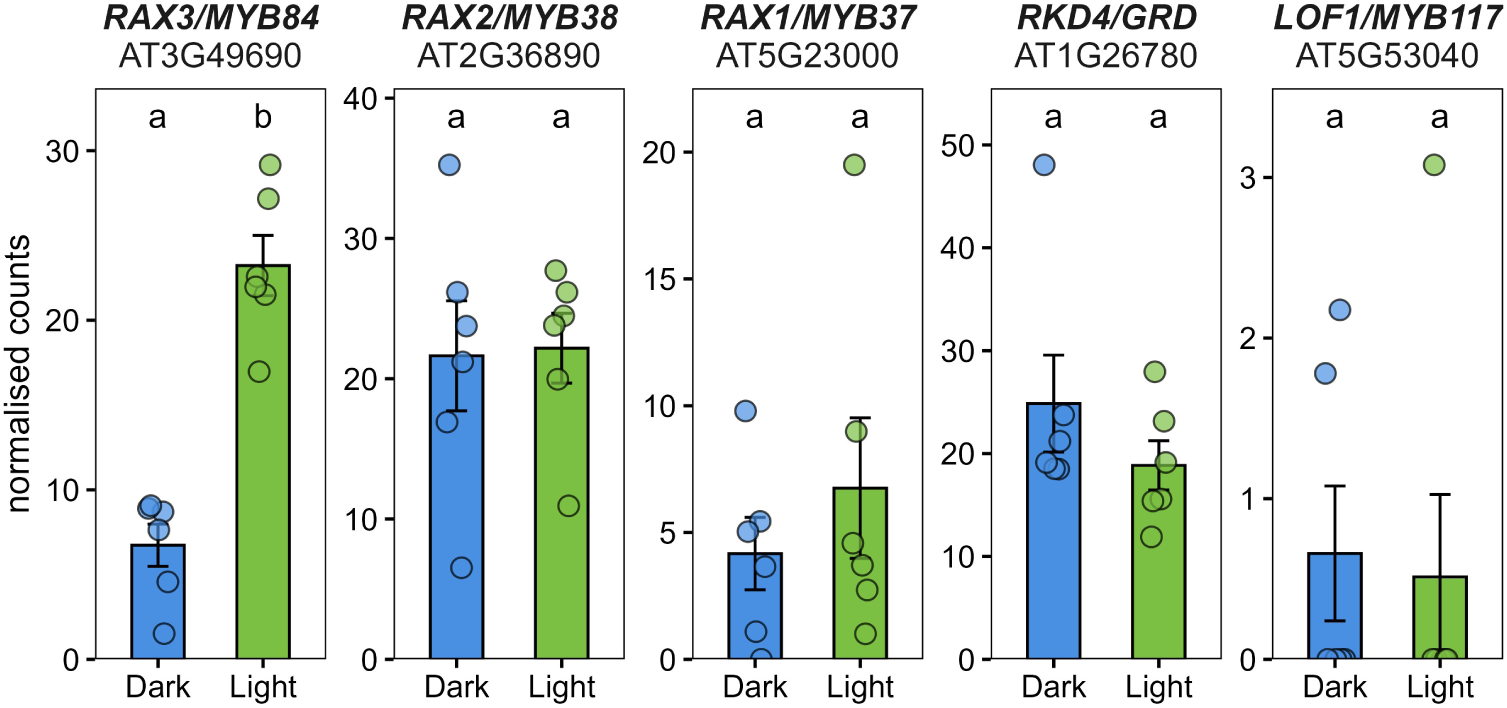
Impact of carbon availability on the expression of gemma cup regulatory gene orthologues in *Arabidopsis thaliana*. Expression of *Arabidopsis thaliana* orthologues of *Marchantia* gemma cup regulatory genes in three-week-old rosettes subjected to normal light (Light) or dark (Dark) conditions. *RAX3/MYB84* is orthologous to *MpGCAM1*; *RAX1/MYB37* and *RAX2/MYB38* are related R2R3-MYB transcription factors; *RKD4/GRD* is orthologous to *MpRKD*; *LOF1/MYB117* is orthologous to *MpSTG*. Data represent mean ± SE of normalised read counts (n = 6 biological replicates, shown as individual points). Different letters indicate statistically significant differences (P < 0.05, DESeq2).

**Supplementary Figure S4.**
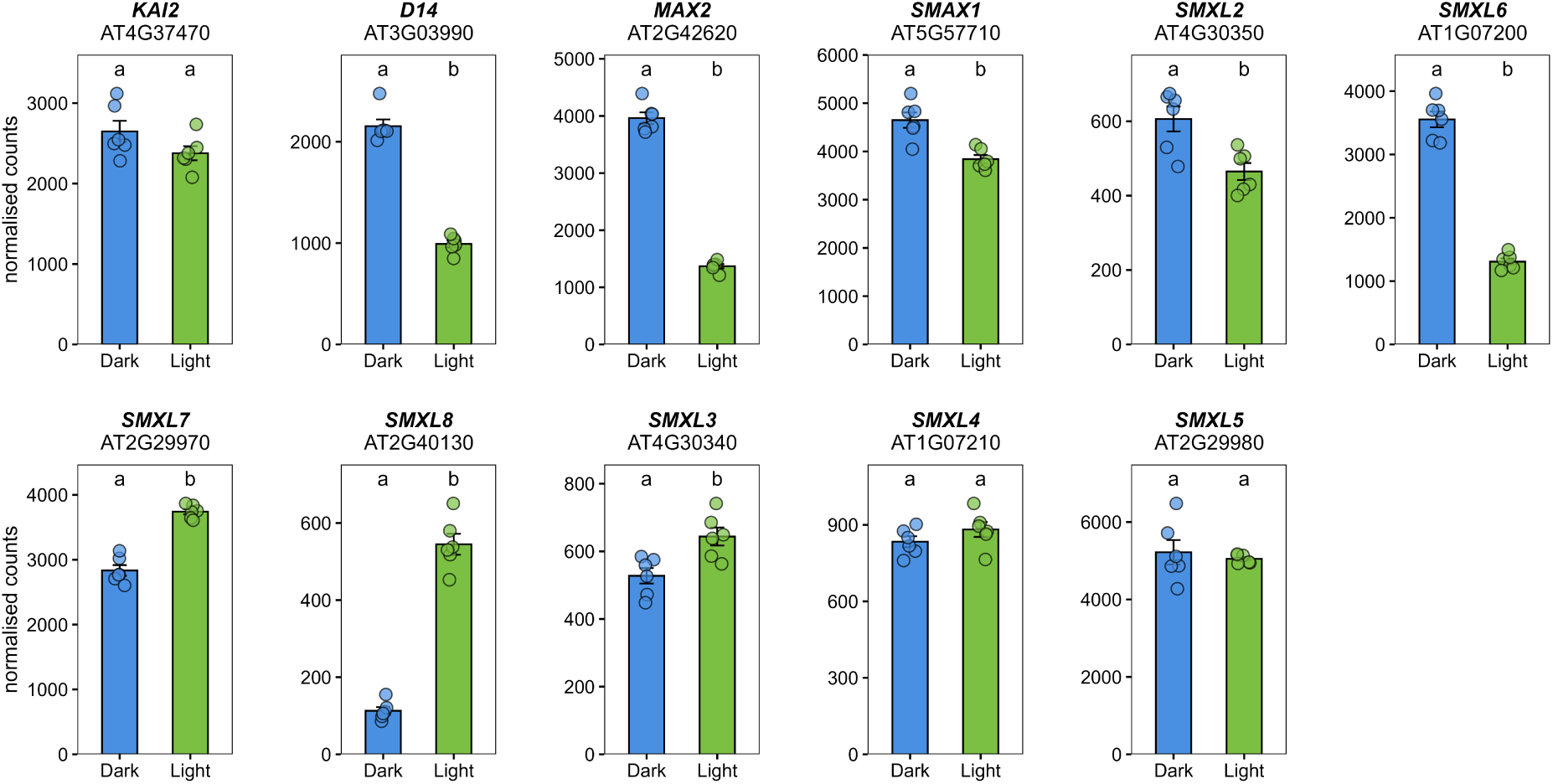
Impact of carbon availability on the expression of *KAI2* and *MAX2* and their downstream targets in *Arabidopsis thaliana*. Expression of *Arabidopsis thaliana* orthologues of genes involved in KAI2- and MAX2-dependent signalling in three-week-old rosettes subjected to normal light (Light) or dark (Dark) conditions. Data represent mean ± SE of normalised read counts (n = 6 biological replicates, shown as individual points). Different letters indicate statistically significant differences (P < 0.05, DESeq2).

